# Imaging specific cellular glycan structures using glycosyltransferases via click chemistry

**DOI:** 10.1101/181453

**Authors:** Zhengliang L Wu, Anthony Person, Matthew Anderson, Barbara Burroughs, Timothy Tatge, Yonglong Zou, Liangchun Wang, Todd Geders, Robert Sackstein

## Abstract

Heparan sulfate (HS) is a polysaccharide fundamentally important for biologically activities. T/Tn antigens are universal carbohydrate cancer markers. Here, we report the specific imaging of these carbohydrates using a mesenchymal stem cell line and human umbilical vein endothelial cells (HUVEC). The staining specificities were demonstrated by comparing imaging of different glycans and validated by either removal of target glycans, which results in loss of signal, or installation of target glycans, which results in gain of signal. As controls, representative key glycans including O-GlcNAc, lactosaminyl glycans and hyaluronan were also imaged. HS staining revealed novel architectural features of the extracellular matrix (ECM) of HUVEC cells. Results from T/Tn antigen staining suggest that O-GalNAcylation is a rate-limiting step for O-glycan synthesis. Overall, these highly specific approaches for HS and T/Tn antigen imaging should greatly facilitate the detection and functional characterization of these biologically important glycans.

## Introduction

Glycosylation is the most common type of post-translational protein modification (Khoury, G.A., Baliban, R.C., et al. 2011). Glycans are assembled by various glycosyltransferases along a pathway starting at the endoplasmic reticulum continuing to the Golgi apparatus, and these structures are expressed on the membrane, or as secreted extracellular matrix elements or soluble glycoproteins (Lairson, L.L., Henrissat, B., et al. 2008). Glycan modifications play various biological roles from protein folding and quality control to a large number of biological recognition events (Moremen, K.W., Tiemeyer, M., et al. 2012). For instance, they act as ligands for numerous glycan-binding lectins (Pang, P.C., Chiu, P.C., et al. 2011), growth factors and cytokines (Dalziel, M., Crispin, M., et al. 2014, Xu, D. and Esko, J.D. 2014). One well-known glycan structure is sialyl Lewis X (sLeX), the canonical binding determinant for selectins, important mediators of cell migration (Somers, W.S., Tang, J., et al. 2000).

Glycans usually are displayed on the cell surface and in the extracellular matrix (ECM) in the forms of glycoproteins (O-glycans and N-glycans), glycolipids (O-glycans) and proteoglycans (glycosaminoglycans)(Bertozzi, C.R., Freeze, H.H., et al. 2009). A representative O-glycan is the “core-1” glycan (Galβ1-3GalNAc-R), also known as “T antigen” that is commonly displayed on various cancer cells (Brockhausen, I., Schachter, H., et al. 2009). Core-1 O-glycan synthesis starts by addition of a GalNAc residue to serine and threonine residues of a protein by a polypeptide GalNAc transferase (GALNT), and is completed by addition of a Gal residue by the β-3 galactosyltransferase C1GalT 1. Loss of the activity of C1GalT 1 in cancer cells results in intermediate product O-GalNAc, also known as “Tn antigen”(Ju, T. and Cummings, R.D. 2005). T and Tn antigens are hallmarks of cancer (Fuster, M.M. and Esko, J.D. 2005, Gilgunn, S., Conroy, P.J., et al. 2013, Pinho, S.S. and Reis, C.A. 2015). Heparan sulfate (HS), a linear polysaccharide that has repetitive disaccharide units of HexA-GlcNAc, is a representative glycosaminoglycan found on the cell surface and in ECM (Iozzo, R.V. 2005, Sarrazin, S., Lamanna, W.C., et al. 2011), with the HexA residue being either a GlcA or IdoA. HS is synthesized by exostosins (EXTs), dual enzymes that have both GlcA and GlcNAc transferase activities (Senay, C., Lind, T., et al. 2000). HS is degraded by heparanase (HPSE), an endoglucuronidase that specifically hydrolyzes the βGlcA-1,4-GlcNAc bond in highly sulfated HS domains(Mao, Y., Huang, Y., et al. 2014, Vlodavsky, I., Friedmann, Y., et al. 1999). HS can also be digested with bacterial lyases such as heparinase III, which leaves an unsaturated GlcA residue (ΔGlcA) at its non-reducing end(Hovingh, P. and Linker, A. 1970). Apart from membrane and extracellular matrix display, proteins within the cytoplasm and nucleus can be modified by O-GlcNAc that plays important regulatory roles in metabolism, diabetes and cancer (Bond, M.R. and Hanover, J.A. 2013, Slawson, C. and Hart, G.W. 2011). O-GlcNAc can be extended with a GalNAc residue by a mutant enzyme B4GalT1Y285L (Ramakrishnan, B. and Qasba, P.K. 2002).

Despite the abundance of glycans and their important biological functions, glycans are notoriously difficult to image due to lack of high affinity and highly specific binding reagents (Sterner, E., Flanagan, N., et al. 2016). In the past decade, metabolic labeling using clickable carbohydrate (Baskin, J.M., Dehnert, K.W., et al. 2010, Codelli, J.A., Baskin, J.M., et al. 2008, Hsu, T.L., Hanson, S.R., et al. 2007, Kizuka, Y., Funayama, S., et al. 2016) has revolutionized glycan imaging, as it eliminates the problems associated with low binding affinity via covalent bonding. Subsequently, enzymatic glycan labeling using clickable sugars have been reported (Griffin, M.E. and Hsieh-Wilson, L.C. 2016): (1) O-GlcNAc has been imaged by further modification using the mutant galactosyltransferase B4GalT1Y285L (Clark, P.M., Dweck, J.F., et al. 2008); (2) the Fucα(1-2)Gal epitope has been imaged by a bacterial glycosyltransferases BgtA (Chaubard, J.L., Krishnamurthy, C., et al. 2012); (3) LacNAc has been imaged by an *H. pylori* fucosyltransferase (Zheng, T., Jiang, H., et al. 2011); and (4) N-glycans have been imaged by recombinant sialyltransferase ST6Gal1 (Mbua, N.E., Li, X., et al. 2013). These methods have shown the promise of specific glycan imaging due to the high specificities of labeling glycosyltransferases. However, the challenge to further prove the specificities of these methods still remains.

Previously, GlcNAc transferases GCNT1, B3GNT6 and EXT1/EXT2 have been found to be permissible to azido-GlcNAc (Wu, Z.L., Huang, X., et al. 2016a, Wu, Z.L., Huang, X., et al. 2016b), and polypeptide GalNAc transferases GALNT1/2/3 have been found to be permissible to azido-GalNAc (Pratt, M.R., Hang, H.C., et al. 2004, Wu, Z.L., Huang, X., et al. 2015). Here, we report the imaging of HS and T/Tn antigens along with other related glycans on mammalian cells by *in vitro* incorporation of azido-sugars using these glycosyltransferases. The incorporated sugars are linked to fluorescent tags via azide-alkyne cycloaddition (Rostovtsev, V.V., Green, L.G., et al. 2002). Using O-GlcNAc as a control, and, HS and T/Tn antigens as examples, the specificities of these methods are systematically investigated through comparing imaging of different glycans, and, imaging before and after removal or installation of the glycans of interests. Leveraging the specificity of these enzymes, the resulting incorporated clickable sugars enable precise detection of the interested glycan motifs and provide key information on the cellular topography of these glycans as well.

## Results

### Glycan imaging on transformed mesenchymal stem cells

C3H10T1/2 cells were originally established from cells extracted from C3H mouse embryos and are an established transformed mesenchymal stem cell model as they can be differentiated into downstream mesenchymal cell lineages(Reznikoff, C.A., Brankow, D.W., et al. 1973). As a proof of concept, we imaged O-GlcNAc, O-GalNAc (Tn antigen), terminal sialyllactosamine (sLN), and core-1 O-glycan (T antigen) using B4GalT1Y285L, B3GNT6, FUT7 and GCNT1, respectively on C3H10T1/2 cells (Table 1). B4GalT1Y285L is a mutant enzyme that can transfer a GalNAc residue to a terminal GlcNAc residue (Ramakrishnan, B. and Qasba, P.K. 2002), and therefore be used for O-GlcNAc detection in the presence of the donor substrate UDP-azido-GalNAc. B3GNT6 specifically recognizes O-GalNAc and synthesizes the “core-3” O-glycan (GlcNAcβ1-3GalNAcα1-S/T) (Iwai, T., Inaba, N., et al. 2002), and is therefore useful for O-GalNAc detection in the presence of UDP-azido-GlcNAc. The α(1,3)-fucosyltransferase FUT7 recognizes sialylated type 2 lactosamine (sLN) and synthesizes sLeX structure (Natsuka, S., Gersten, K.M., et al. 1994), and is thus useful for sLN detection in the presence of GDP-azido-fucose. GCNT1 transfers a GlcNAc residue to core-1 O-glycan and synthesizes “core-2” O-glycans (Yeh, J.C., Ong, E., et al. 1999), and can thereby be used to detect core-1 O-glycan in the presence of UDP-azido-GlcNAc.

**Table I.**
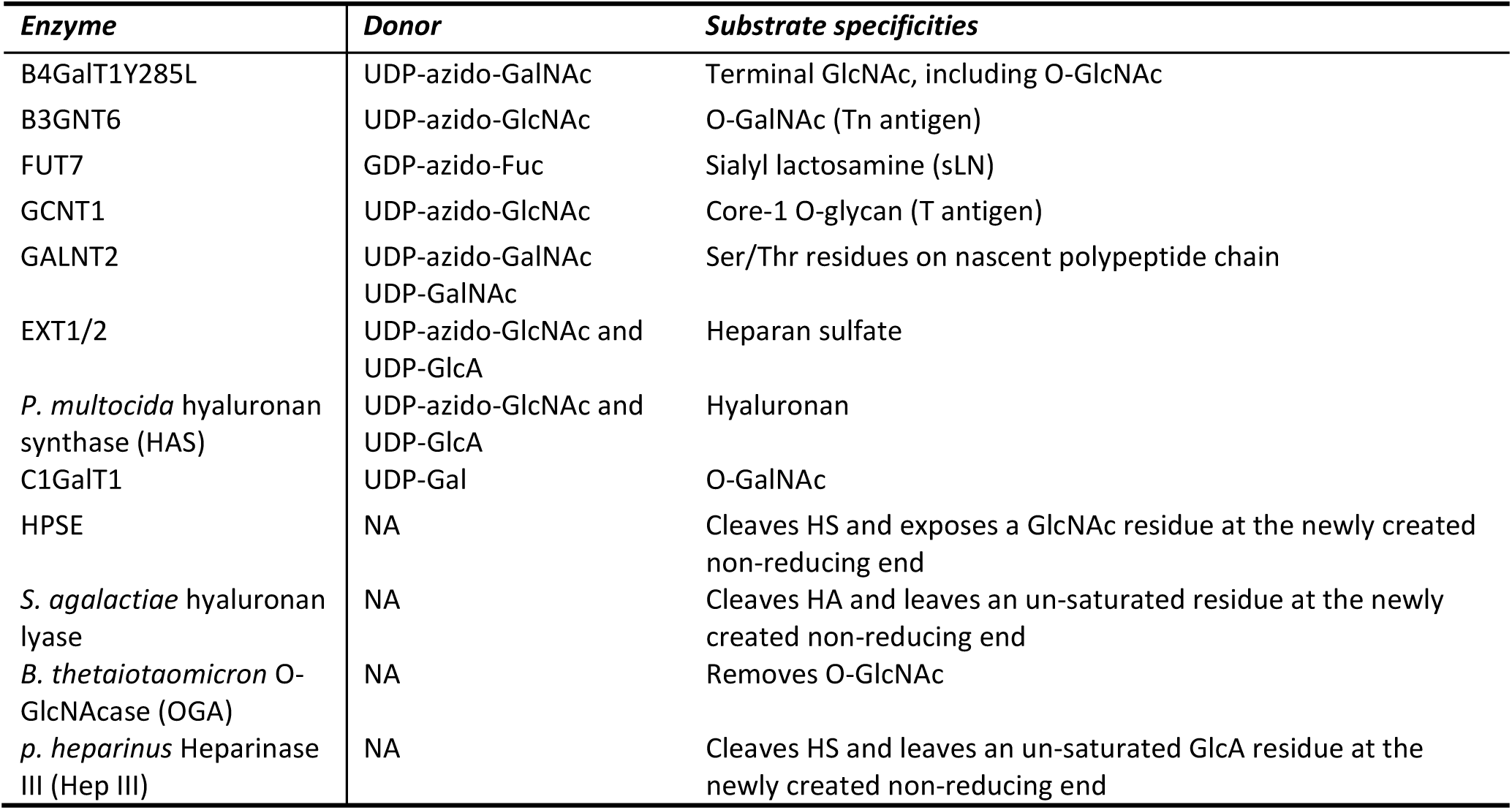
Enzymes used in this study. All are recombinant human enzymes except indicated otherwise.

Consistent with results of others (Hart, G.W. and Akimoto, Y. 2009, Wu, Z.L., Robey, M.T., et al. 2014), B4GalT1Y285L stained strongly in the nuclei and weakly in regions surrounding the nuclei (Fig. 1A). Direct staining for O-GalNAc (Tn antigen) using B3GNT6 resulted in no signal. To assess whether Tn antigen can be detected, it was first installed on cells by GALNT2, a polypeptide GalNAc transferase that transfers a GalNAc residue to serine/threonine residues on nascent polypeptides (Sorensen, T., White, T., et al. 1995), and then probed using B3GNT6. The installed O-GalNAc was mainly found in the cytoplasm (Fig. 1B), which is reasonable because the labeled proteins are likely to be nascent polypeptides that contained unmodified Ser/Thr residues. In sharp contrast, FUT7 and GCNT1 labeling appeared to be largely on thecell membrane and the ECM (Fig. 1C and 1D, respectively). The resultant staining using FUT7 indicates that there are sLN on C3H10T1/2 cells, which is consistent with previous findings that both mouse and human mesenchymal stem cells can be α(1,3)-exofucosylated to enforce sLeX expression (Abdi, R., Moore, R., et al. 2015, Dykstra, B., Lee, J., et al. 2016). The staining observed using GCNT1 indicates that C3H10T1/2 cells express core-1 O-glycan or T antigen.

**Figure 1.**
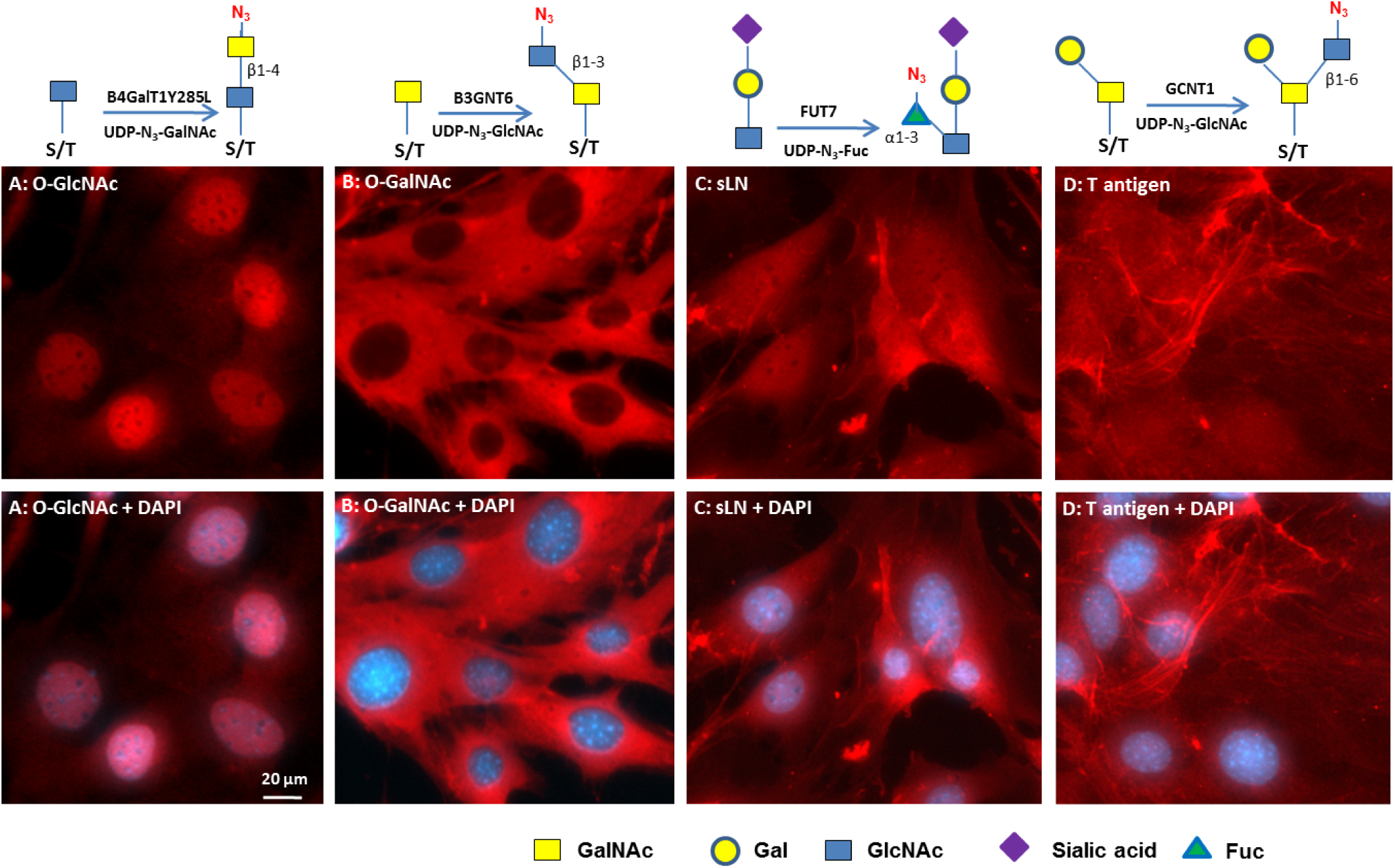
Visualizing glycans on mesenchymal C3H10T1/2 cells. The cells grown to confluence were imaged for the indicated glycans using corresponding glycosyltransferases. The glycans were revealed with Alexa-Fluor 555 (red). Nuclei were revealed with DAPI (blue). All images were normalized to their highest pixel value without change of gamma rating. Labeling strategies are indicted above the pictures. **A)** O-GlcNAc imaging. **B)** O-GalNAc imaging. O-GalNAc was installed by GALNT2 in the presence of UDP-GalNAc. **C)** Sialyllactosamine (sLN) imaging. **D)** Core-1 O-glycan imaging.

### Glycan imaging on human umbilical vein endothelial cells

Human umbilical vein endothelial cells (HUVEC) are primary endothelial cells commonly used to study the mechanisms of angiogenesis *in vitro* (Crampton, S.P., Davis, J., et al. 2007). HUVEC cells were grown in a 24-well plate to confluence and then stained for O-GlcNAc, O-GalNAc, sLN, HS and HA determinants using recombinant human B4GalT1Y285L, B3GNT6, FUT7 and EXT1/2, and recombinant *Pasteurella multocida* HA synthase (HAS) respectively (Table 1). EXT1/2 is a hetero-dimeric HS polymerase of EXT1 and EXT2, which alternatively utilizes UDP-GlcA and UDP-GlcNAc to extend the HS chain *in vivo.* Previously, we showed that EXT1/2 can incorporate azido-GlcNAc to the non-reducing ends of HS (Wu, Z.L., Huang, X., et al. 2016b). HAS is a due polymerase that can alternatively utilizes UDP-GlcA and UDP-GlcNAc to extend an HA chain at its non-reducing end (DeAngelis, P.L., Oatman, L.C., et al. 2003).

Similar to C3H10T1/2 cells, O-GlcNAc staining was primarily found in the nuclei and secondarily found in regions close to the nuclei (Fig. 2A). Again, “free” O-GalNAc (Tn antigen) was not identifiable in HUVEC cells so it was installed by GALNT2 prior to B3GNT6 staining, and similar to the C3H10T1/2 cells, the incorporated O-GalNAc was primarily located in the cytoplasm (Fig. 2B). FUT7 staining yielded strong signal on the cell body and weak signal within the ECM (Fig. 2C); suggesting a higher density of sLN on the cell surface than in the ECM. In contrast, EXT1/2 yielded strong staining of the ECM but weak staining of the cell surface (Fig. 2D, see also Figure S1). In fact, the staining of cell body by EXT1/2 was so weak that the cell bodies were only revealed by GALNT2 staining in the presence of UDP-azido-GalNAc in Fig. 2D.

**Figure 2.**
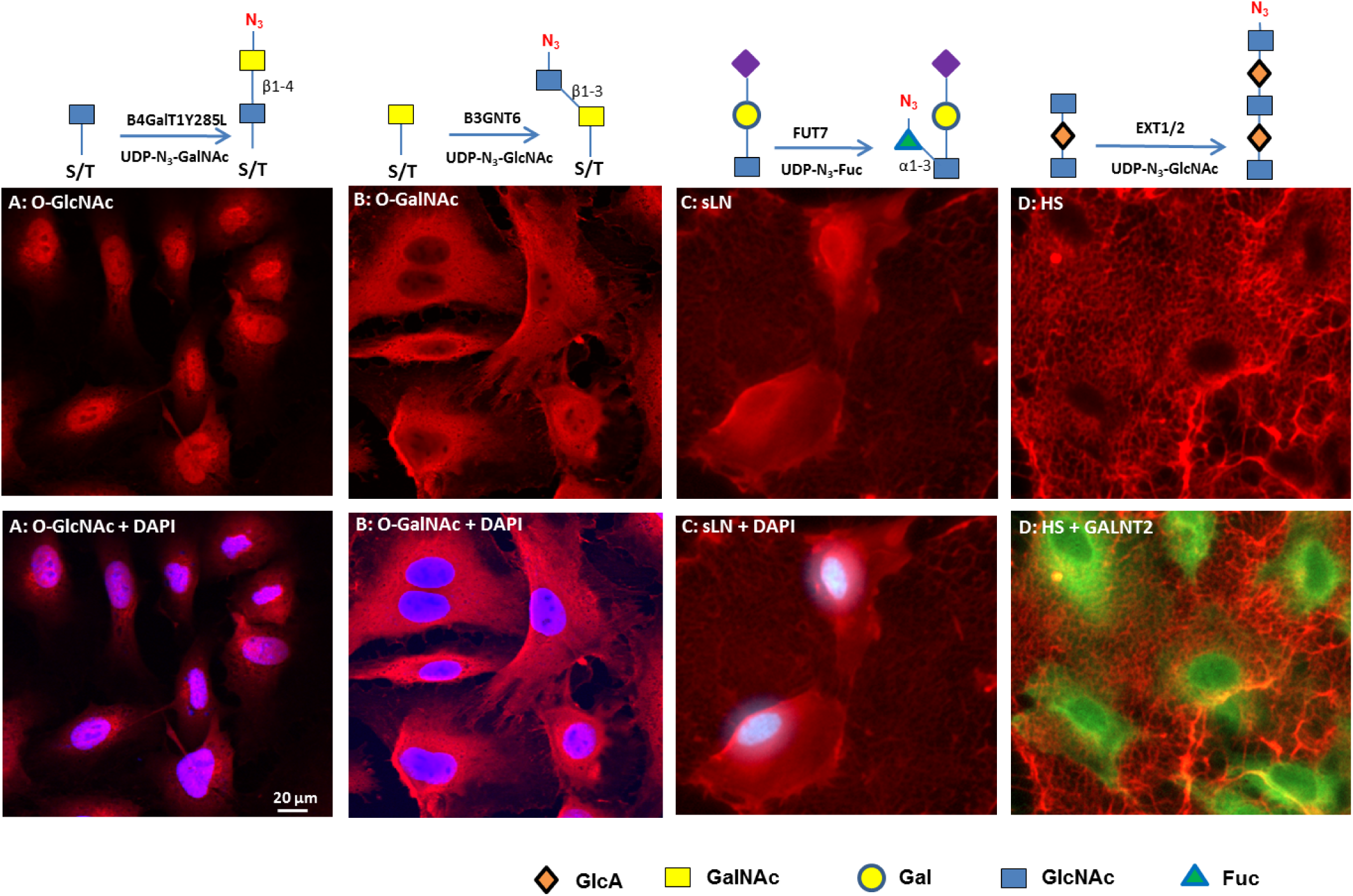
Visualizing glycans on HUVEC cells. HUVEC cells were grown in a 24-well plate to sub-confluence. The indicated glycans were then imaged using corresponding enzymes. The glycans were revealed with Alexa-Fluor 555 (red). Nuclei were revealed with DAPI (blue). All images were normalized to their highest pixel value without change of gamma rating. **A)** O-GlcNAc imaging. **B)** O-GalNAc imaging. O-GalNAc was installed by GALNT2 in the presence of UDP-GalNAc. **C)** Sialyllactosamine (sLN) imaging. **D)** HS imaging. To increase HS staining, cells were pretreated with HPSE. In the lower panel of **D**, cells were also imaged using GALNT2 /UDP-azido-GalNAc and Alexa-Fluor 488 (green).

HUVEC cells were also stained for HA (Figure S2). While HS was located mainly in ECM and weakly detected on cell body, HA was strictly found on cell body and particularly enriched in nucleus. The HA detected on cell body and nucleus could be those molecules that were bound to their receptors such as CD44 (Aruffo, A., Stamenkovic, I., et al. 1990). In a separate experiment, prior treatment of HUVEC cells with HA lyase abolished HA staining (Figure S3), confirming the identity of HA that was detected.

### Further confirmation of the specificity of glycan imaging

From the above examples of cell staining, it is clear that different enzymes stained different features on cells, indicating that this glycosyltransferase-mediated staining is specific. To further assess whether staining was indeed on the target glycans that are recognized by the enzymes, we utilized two strategies. The first involved abolishing staining by removal of target glycans using specific glycosidases, which is already shown in HA staining (Figure S3). The second involved increased staining by installation of target glycans using glycosyltransferases. In some cases, the results of cell staining were further supported by results obtained from labeling of cellular extracts detected by the GLCC method previously described (Wu, Z.L., Huang, X., et al. 2015).

### Specificity of O-GlcNAc staining

Since O-GlcNAc is well known for its cellular localization (Hart, G.W. and Akimoto, Y. 2009), it was chosen for testing the specificity of the current method of imaging. For this purpose, we treated HUVEC cells with O-GlcNAcase (OGA), a glycosidase that specifically removes O-GlcNAc (Dennis, R.J., Taylor, E.J., et al. 2006) but not terminal GlcNAc, prior to the staining with B4GalT1Y285L (Figure 3). As expected, the nuclear staining was abolished after the treatment, which confirms that the staining in the nuclei is on O-GlcNAc. The remaining staining in the regions surrounding the nuclei is likely on terminal GlcNAc residues other than O-GlcNAc, as it is known that the mutant B4GalT1 also uses terminal GlcNAc residues on N-glycans and O-glycans as acceptor substrates (Boeggeman, E., Ramakrishnan, B., et al. 2007).

**Figure 3.**
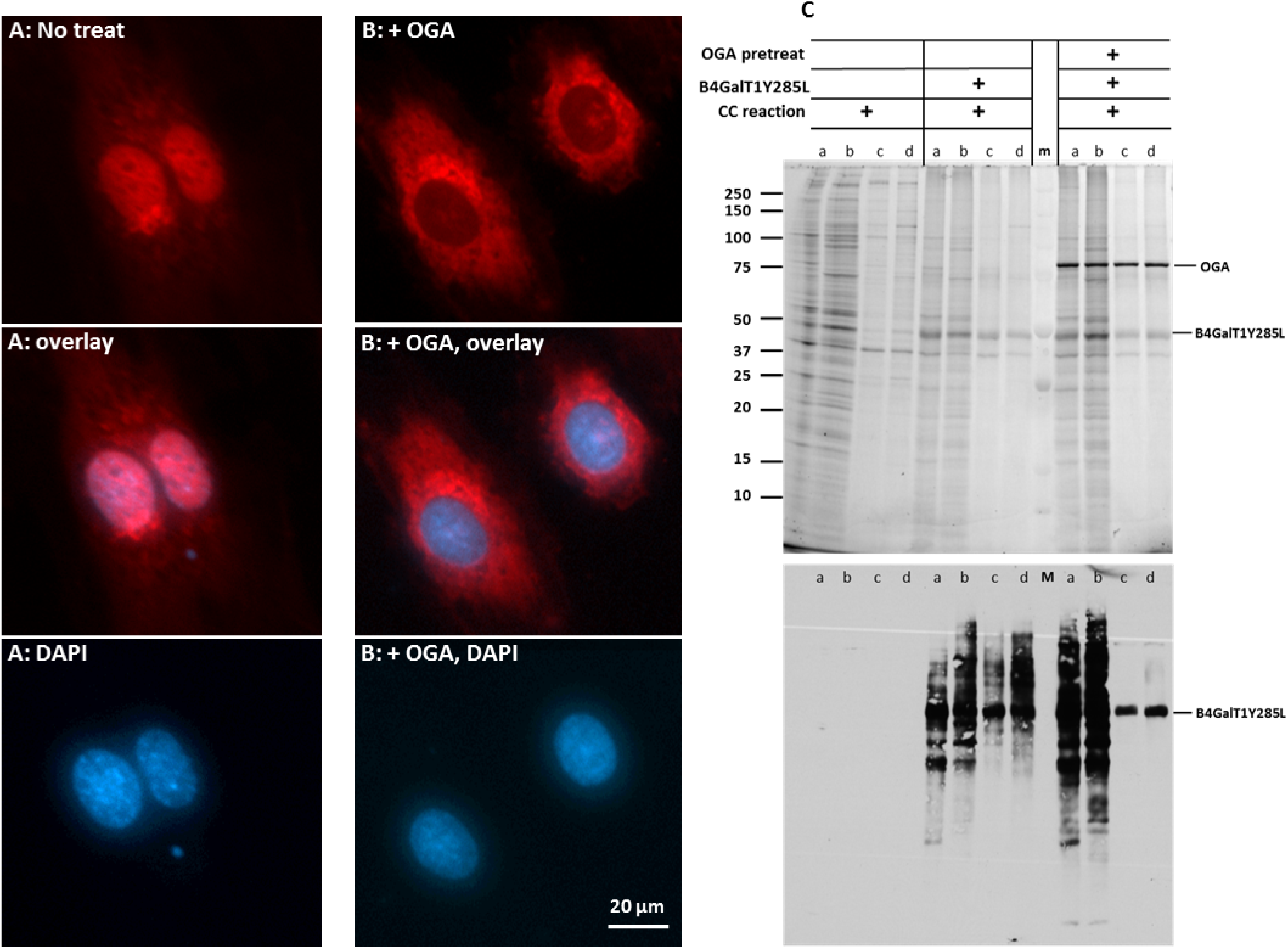
Specificity of O-GlcNAc imaging. HUVEC cells were grown in a 24-well plate to sub-confluence and then imaged directly using B4GalT1Y285L (A), or imaged after the pretreatment of OGA (B). O-GlcNAc was stained in red and nuclei were in blue. All images of O-GlcNAc staining were normalized to the highest pixel value without changing the gamma setting. (C) Detection of O-GlcNAc in cytoplasmic and nuclear extracts of HUVEC and HEK 293 cells. The extracts were labeled for O-GlcNAc by B4GalT1Y285L either directly or after pretreatment of OGA. All samples were subject to click chemistry (CC) reaction Upper panel, SDS-PAGE of the samples. Lower panel, blotting of the upper panel with Streptavidin-HRP. a, cytoplasmic extract from HUVEC cell; b, cytoplasmic extract from HEK 293 cells; c, nuclear extract from HUVEC cells; d, nuclear extract from HEK cells; m, pre-stained molecular weight marker.

In a parallel experiment, cytoplasmic and nuclear extracts prepared from HUVEC and HEK 293 cells were treated with OGA first and then labeled with B4GalT1Y285L (Figure 3C). Both cytoplasmic and nuclear extracts were labeled in the absence of OGA treatment. In contrast, OGA abolished the labeling of nuclear extracts only, again confirming that the labeling in the nuclear extracts was exclusively on O-GlcNAc.

### Strict specificity of HS staining

To test the specificity of HS staining, we treated HUVEC cells with heparinase III (Hep III) and heparanse (HPSE), prior to the staining by EXT1/2 (Fig. 4). Hep III digestion on HS results in HS chains ending with Δ4,5-glucuronic acid (ΔGlcA) residues that lacks a C4-OH group that is needed for HS chain extension by EXT1/2. Indeed, Hep III digestion abolished the labeling by EXT1/2 (Fig. 4B), confirming that the staining is indeed on HS. On the other hand, HPSE digestion on HS results in uniform N-sulfated GlcNAc residue (GlcNS) at the ends of HS chains(Mao, Y., Huang, Y., et al. 2014). As a result, pretreatment with HPSE significantly increased the labeling (Fig. 4C), suggesting that GlcNS can be extended by EXT1/2 and that a significant portion of native HS chains could not be labeled by EXT1/2. Finally, we attempted to label HPSE treated cells without the donor substrate UDP-GlcA. This experiment resulted in almost no labeling in the ECM (Fig. 4D), again confirming that the labeling was on HS, because HS digested by HPSE ends with a GlcNS residue(Mao, Y., Huang, Y., et al. 2014) that requires the incorporation of a GlcA residue prior to the incorporation of an azido-GlcNAc for HS detection.

**Figure 4.**
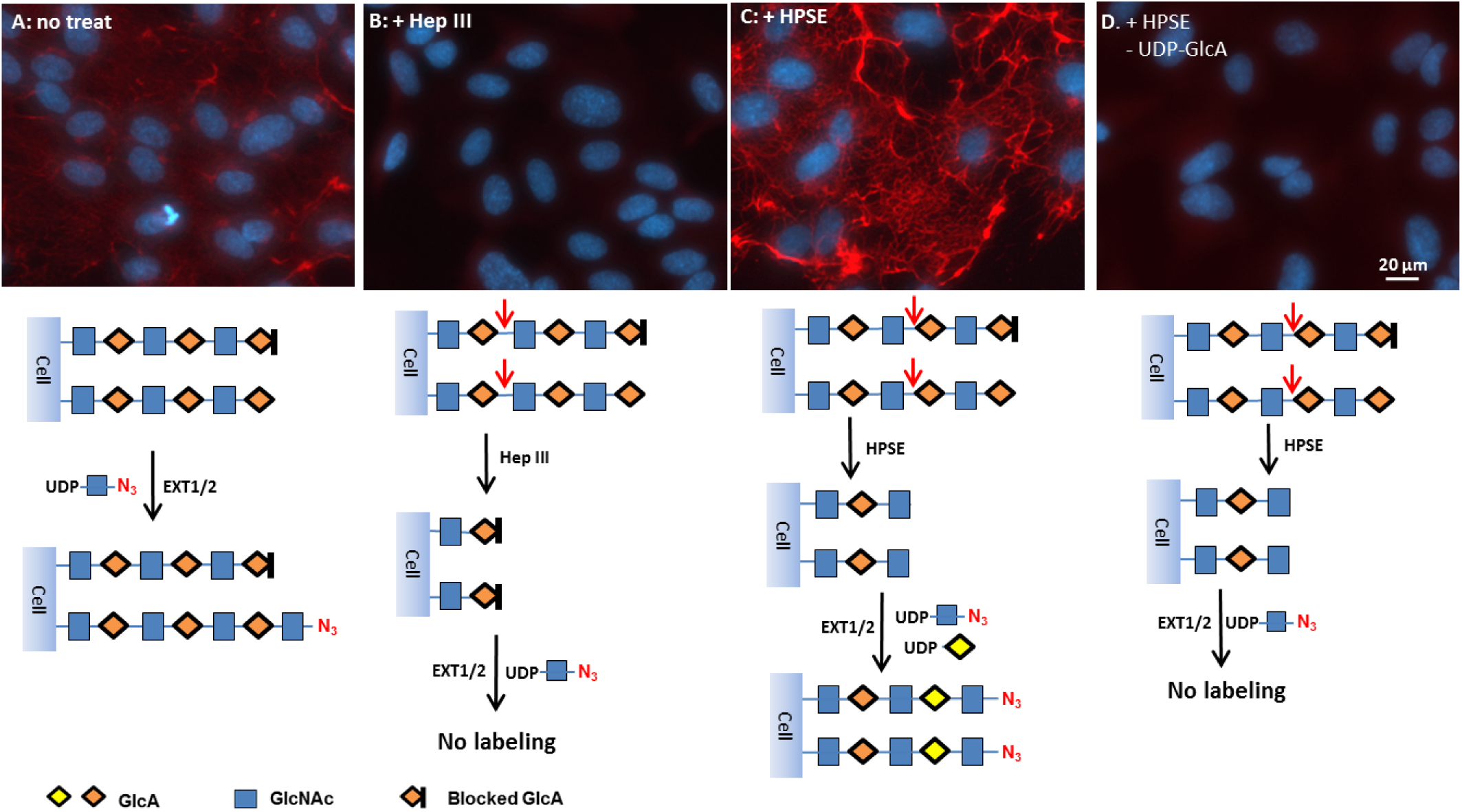
Specificity of heparan sulfate imaging. HUVEC cells were grown in a 24-well plate to confluence and then HS was imaged directly using EXT1/2 **(A)**, imaged after the pretreatment of Hep III **(B)**, imaged after the pretreatment of HPSE with both UDP-azido-GlcNAc and UDP-GlcA **(C)**, imaged after the pretreatment of HPSE with UDP-azido-GlcNAc only. HS was stained in red and nuclei were revealed with DAPI (blue). For comparison, exposure time, contrast and gamma setting were kept the same for all the pictures. Schemes for the labeling strategies are showing below each panel.

### ECM morphology revealed by HS staining

While HS was found mainly in the ECM of confluent HUVEC cells (Fig. 2D), it was also found on cells (Figure S1 and Figure S2), which is consistent with the literature (Sarrazin, S., Lamanna, W.C., et al. 2011). HS staining also revealed some unique architecture of the ECM of HUVEC cells. In one case, the ECM resembled an entangled mesh (Figure 5A). In another case, the ECM resembled a lush lawn (Figure 5B). In a third case, the ECM resembled a plain carpet (Figure 5C), suggesting the cells that had laid out the HS had migrated away from sites ofdeposition. The unique architecture of the HS in each case suggests that dynamic alterations in HS topography may dictate cellular distribution in discrete tissue microenvironments. When HS of non-confluent young HUVEC cells was stained, some intermediate structures of ECM were captured (Figure S4), suggesting that the ECM formation by HUVEC cells may follow a unique mechanism.

**Figure 5.**
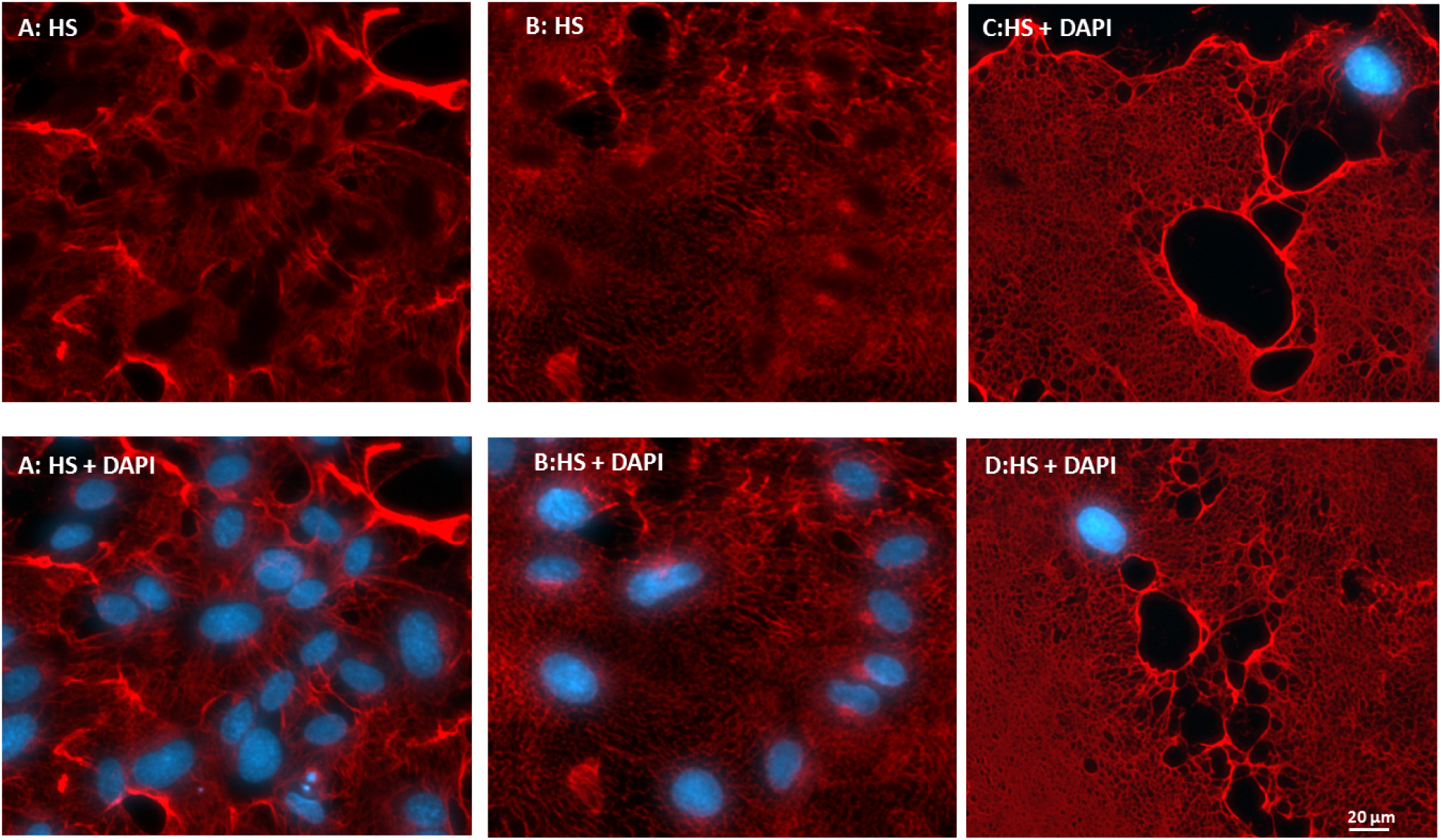
Morphology of ECM of HUVEC revealed by HS staining. HUVEC cells grown in a 24-well plate to confluence were treated with HPSE and followed by HS imaging using EXT1/2 with UDP-azido-GlcNAc and UDP-GlcA. HS imaging was revealed by Alexa-Fluor 555 (red) and nuclei were revealed with DAPI (blue). The staining on ECM was so strong that the staining on the cell body was not visible. (A), the ECM resembles a tangled mesh. (B), the ECM resembles a lush lawn. (C), the ECM resembles a plain carpet.

### Enzymatic synthesis of T and Tn antigens and their specific staining

To assess whether GCNT1 staining observed in Fig. 1D is indeed on T antigen (or core-1 O-glycan), we installed T antigen to HUVEC cells that natively do not express the glycan using GALNT2 and C1GalT1. GALNT2 attaches a GalNAc residue to serine or threonine residue on the protein backbone to generate O-GalNAc(Sorensen, T., White, T., et al. 1995), and C1GalT1 further adds a Gal residue to the O-GalNAc to complete the core-1 O-glycan (T antigen) synthesis(Ju, T., Brewer, K., et al. 2002) (Fig. 6A). When the T antigen was fully installed, the staining with GCNT1 was positive (Fig. 6B); while when only Tn antigen was installed, the cell staining by GCNT1 was negative (Fig. 6C). The installation of Tn antigen was confirmed by B3GNT6 staining in Fig. 6D. In contrast, only weak background was seen on untreated cells by B3GNT6 staining (Fig. 6E). These experiments demonstrate the strict specificities of GCNT1 and B3GNT6, for T and Tn antigens, respectively. It should be noted that some staining in the ECM by GCNT1 was observed regardless of T antigen installation (Indicated with arrows in Fig. 6B and 6C), suggesting that HUVEC cells secreted T antigen to the ECM despite the fact that these cells did not display this determinant. In comparison, C3H10T1/2 cells not only secreted T antigen but also displayed this glycan on cell surface (Fig. 1D).

**Figure 6.**
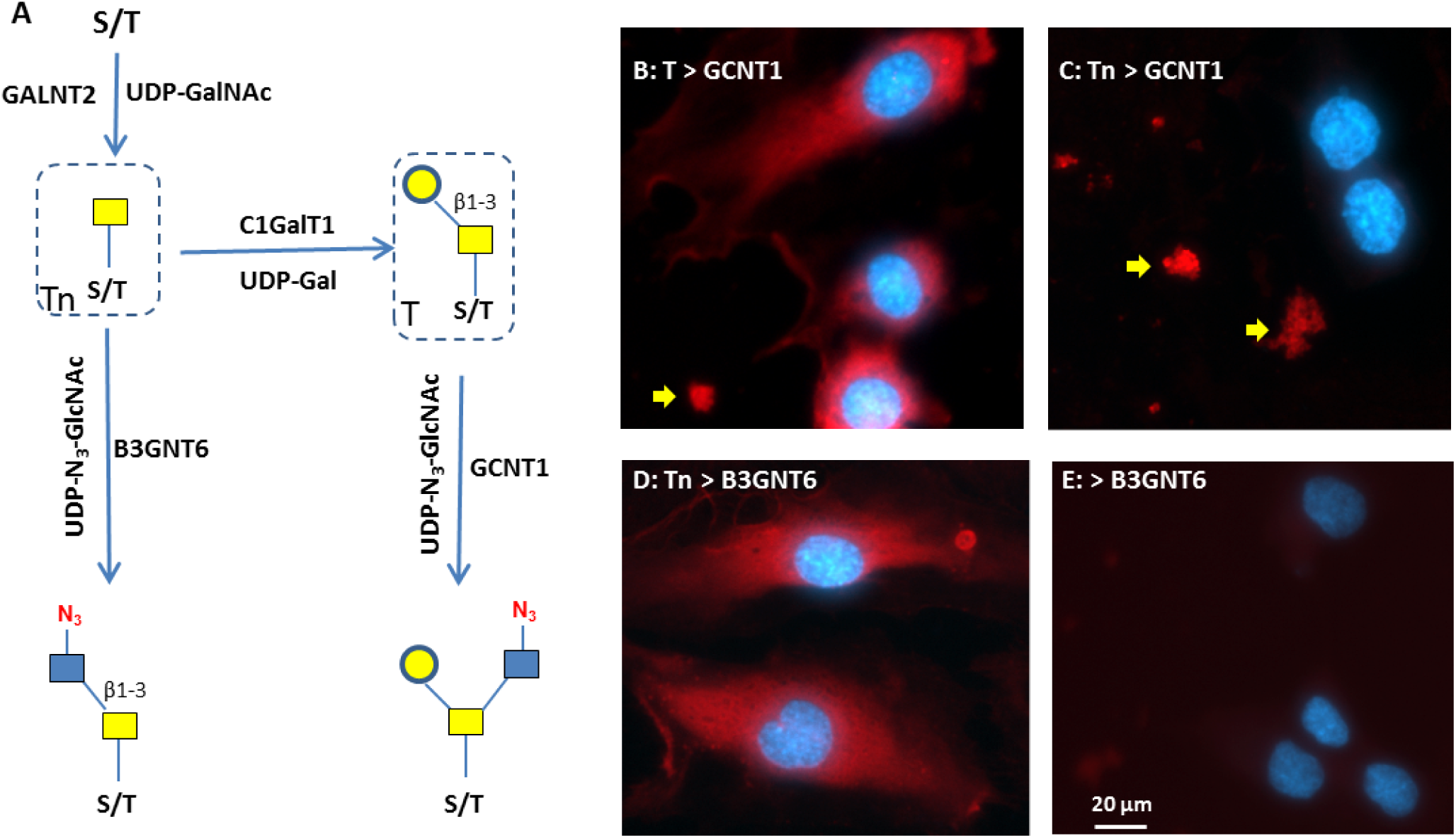
Specificity of T and Tn antigens imaging. **(A)** Schematic view of the synthesis of T and Tn antigens and their detection. The antigens are installed to proteins using GALNT2 and C1GalT1 in the presence of donor substrates UDP-GalNAc and UDP-Gal. T antigen is detected by GCNT1. Tn antigen is detected by B3GNT6. **(B)** Fully installed T antigen (core-1 O-glycan) on HUVEC cells imaged by GCNT1. **(C)** In the absence of UDP-Gal, only Tn antigen was synthesized, which resulted in negative cell staining by GCNT1. **(D)** In the absence of UDP-Gal, only Tn antigen was synthesized, which resulted in positive cell staining by B3GNT6. **(E)** Fixed HUVEC cells were directly stained by B3GNT6. Staining in **B** and **C** indicated by the yellow arrows suggests the secretion of T antigen by HUVEC cells regardless of the installation. Antigen staining was revealed by Alexa-Fluor 555 (red) and nuclei were revealed by DAPI (blue).

Using the same strategy of Fig. 6A, cytoplasmic and nuclear extracts of HUVEC cells and HEK293 cells were installed with Tn and T antigens and then labeled with B3GNT6 and GCNT1. As expected, only Tn antigen was detected by B3GNT6 (Fig. 7A) and only T antigen was detected by GCNT1 (Fig. 7B). Also, in sharp contrast to B4GalT1Y285L that can label the unmodified cellular extracts (Figure 3C), no labeling was observed with either GCNT1 or B3GNT6 on unmodified cellular extracts, suggesting that these cellular extracts were devoid of T and Tn antigens. These results further provided the evidence of the strict specificities of the labeling enzymes of GCNT1 and B3GNT6. The lack of T and T n antigens on HUVEC cells andthe fact that O-GalNAc can be installed onto fixed HUVEC cells by GALNT2 also suggest that O-GalNAcylation is a rate limiting step for O-glycan synthesis.

**Figure 7.**
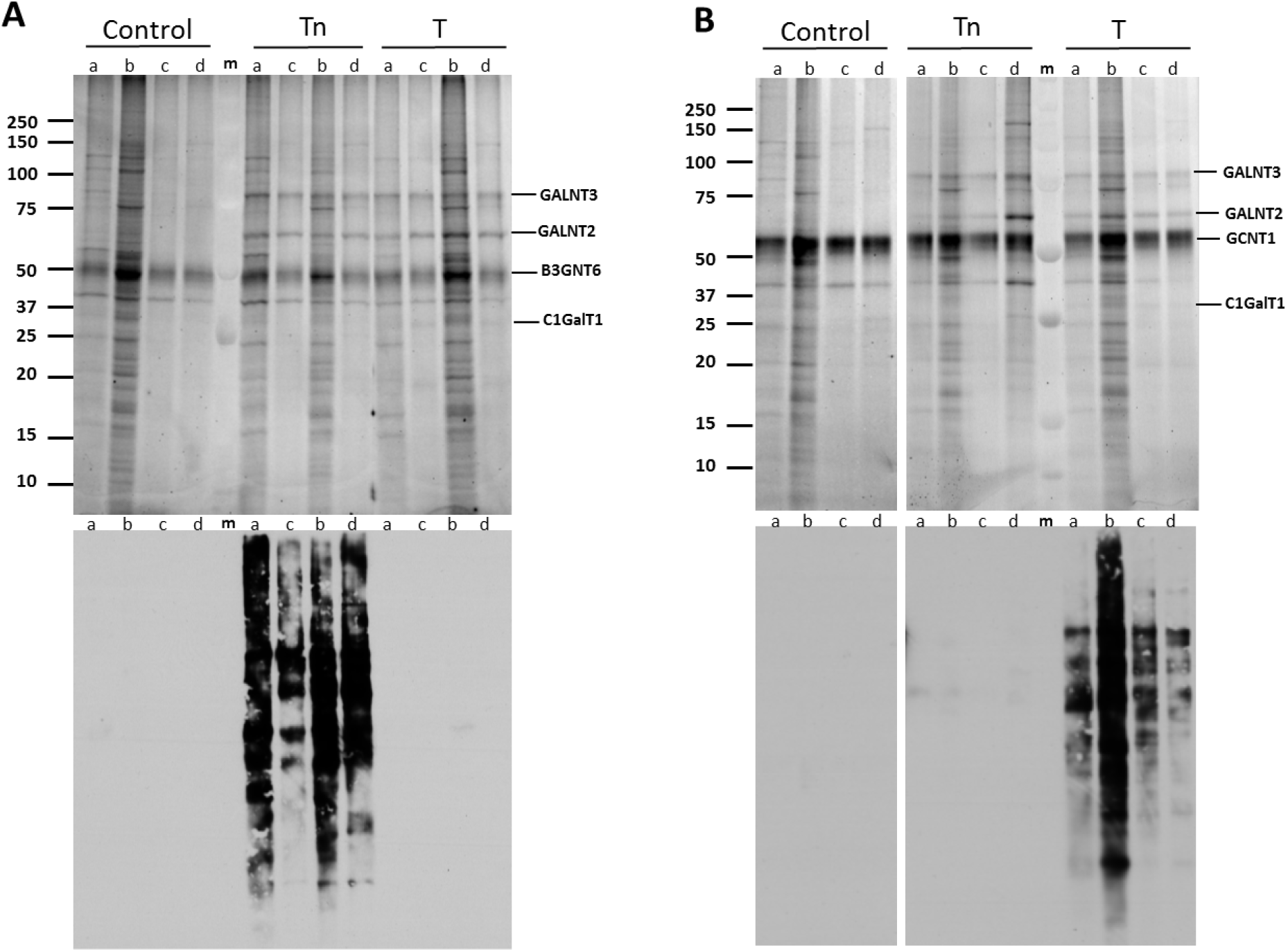
Specific detection of T and Tn antigens installed on cellular extracts. Tn or T antigens were installed to the nuclear and cytoplasmic extracts of HUVEC and HEK 293 cells and then detected with B3GNT6 and GCNT1, using the same strategy as Fig. 5A. **(A)** Detection of Tn antigen installed on the cell extracts by B3GNT6. **(B)** Detection of T antigen installed on cell extracts by GCNT1. Upper panels are SDS-PAGE of the samples, and lower panels are blotting with Streptavidin-HRP. a, cytoplasmic extract from HUVEC cells; b, cytoplasmic extract from HEK 293 cells; c, nuclear extract from HUVEC cells; d, nuclear extract from HEK cells; m, pre-stained molecular weight marker.

Since there are about 20 GALNTs that can perform O-GalNAcylation (Gerken, T.A., Jamison, O., et al. 2011), the substrate specificities and kinetics of these enzymes will be crucial for the synthesis of O-glycans. To this end, we labeled the cellular extracts of HUVEC cells using GALNT1, 2, 3 with UDP-azido-GalNAc (Figure S5). The three enzymes exhibited different labeling patterns, confirming their different substrate specificities (Pratt, M.R., Hang, H.C., et al. 2004). Moreover, the three enzymes showed synergistic effects on O-GalNAcylation. This experiment suggests that for the synthesis of T and Tn antigen on a particular polypeptide scaffold, the selection of GALNT is critical.

## Discussion

Currently, antibodies and plant lectins are the most commonly used reagents to detect glycans (Cummings, R.D. and Etzler, M.E. 2009). However, there are relatively few anti-glycan antibodies and their specificities are generally not absolute (Sterner, E., Flanagan, N., et al. 2016). Though many lectins are characterized, these reagents bind with low affinity to their glycan targets and, thus, generally lack both sensitivity and specificity (Ambrosi, M., Cameron, N.R., et al. 2005). Here, we describe imaging methods especially for heparan sulfate and T/Tn antigen detection by *in vitro* enzymatic incorporation of clickable monosaccharides. The specificity of our methods is achieved through glycosyltransferases that are known to be highly selective for their substrate glycans. By covalent conjugating reporter molecules to target molecules, the sensitivity of the current methods is maximized in theory. The strict specificity of these methods was further confirmed by either removal of target glycans, which resulted in loss of signal, or by installation of a target glycan, which resulted in gain of signal. The images obtained with the described methods either confirm what we already know about a particular glycan or stimulate new questions about the biology of these glycans. In particular, HS staining revealed striking images of the ECM of HUVEC cells, suggesting unique mechanism for HS secretion. T/Tn antigen staining on HUVEC suggested that O-GalNAcylation is a rate limiting step for O-glycan synthesis. We believe that these new tools for glycan imaging will greatly facilitate us to understand the biological roles of various glycans and glycoconjugates.

T and Tn antigens are hall marks of many types of cancer and they become the targets for cancer immunotherapy (Posey, A.D., Jr., Schwab, R.D., et al. 2016). Here we have provided pure enzymatic ways for their synthesis on cellular scaffold as well as methods for their detection. This enzymatic synthetic approach provides an alternative way for enforcing T and Tn antigens, which may allow for further elucidation of the roles of these glycans in cancer biology. It is also foreseeable that the unprecedented specificities of the labeling enzymes for T/Tn antigen maybe exploited for therapeutic interventions such as for linking various bioactive molecules to target cells. Further studies using these reagents should help unveil how dynamic changes in glycan display mediate cellular processes, such as cell-cell and cell-matrix interactions.

## Materials and methods

GDP-azido-Fucose, UDP-azido-GalNAc, UDP-azido-GlcNAc, Biotinylated Alkyne, and 4’,6-diamidino-2-phenylindole (DAPI) were from R&D Systems/Bio-techne. Recombinant human B4GalT1Y285L, GALNT2, FUT7, GCNT1, B3GNT6, EXT1/2, ST6Gal1, C1GalT1 and HPSE, and recombinant *B. thetaiotaomicmn* O-GlcNAcase (OGA), *P.heparinus* heparinase III (Hep III), *P. multocida* hyaluronan synthase (HAS) and S. agalactiae hyaluronan lyase were also from R&D System/Bio-techne. Streptavidin-Alexa Fuor 555, streptavidin-Alexa Fluor 488 and Cy5-Alkyne were from ThermoFisher Scientific. UDP-Gal and UDP-GlcA and all other small chemicals were from Sigma Aldrich.

### Cell growth and fixation

C3H10T1/2 cells (ATCC # CCL-226) were grown in MEM NEAA Earle’s Salts (Irvine Scientific, #9130), supplemented with 10% fetal bovine serum (Corning, #35-015-CV), 2 mM L-glutamine, 100 units/ml penicillin and 0.1 mg/ml streptomycin (Sigma, #G6784). HUVEC (Clonetics/Lonza, #C2517A) were grown in EGM-2 Bullet Kit (Clonetics/Lonza, #CC-3162). Upon confluence, cells were trypsinized and plated in a 24-well cell culture plate and grown to desired confluence. The cells were rinsed with sterile phosphate buffered saline (PBS) and fixed in 4% paraformaldehyde for 30 minutes at room temperature followed by washing 5 times with sterile PBS. After washing, the plate was stored in one milliliter sterile PBS at 4 °C until ready for glycan labeling.

### Pretreatment of cells for imaging

For glycan imaging, the cells were pretreated with glycosidases to remove existing glycans or with glycosyltransferases to add glycans. All of the following treatments are referring to the cells in a single well in a 24-well plate. In particular, for O-GlcNAc removal, cells were treated with 2 μg of OGA in 200 μ of 50 mM MES and 100 mM NaCl at pH 5.5 for 1 hour at 37 °C. For heparan sulfate removal, cells were treated with 2 μg of Hep III in 200 μ of 50 mM Tris and 2 mM CaCl_2_ at pH 7.5 for 1 hour at 37 °C. For heparan sulfate treatment by HPSE, cells were digested by 2 μg of HPSE in 200 μl of 0.1 M NaOAc at pH 4.0 for 1 hour at 37 °C. For installation of O-GalNAc to cells, 50 nmol of UDP-GalNAc and 2 μg of GALNT2 in 200 μl of 25 mMTris, 150 mM NaCl, 10 mM MnCl_2_ at pH 7.5 were added into each well and the cells were then incubated at 37 °C for one hour. After the treatment, the cells were thoroughly washed three times with PBS and all solutions were removed by aspiration.

### Incorporation of clickable carbohydrate using glycosyltransferases

To incorporate clickable carbohydrates into the cells, 25 mM MES, 0.5% (w/v) Triton^^®^^ X-100, 2.5 mM MgCl_2_, 10 mM MnCl_2_, 1.25 mM CaCl_2_ and 0.75 mg/mL of BSA at pH 7.0 was used as the Labeling Buffer throughout the course of this study. For imaging lactosamine, a mixture of 20 nmol of GDP-azido-fucose and 2 μg of FUT7 in 200 μl Labeling Buffer was prepared. For O-GlcNAc imaging, a mixture of 20 nmol of UDP-azido-GalNAc and 2 μg of B4GalT1Y285L in 200 μl Labeling Buffer was prepared. For O-GalNAc imaging, a mixture of 20 nmol of UDP-azido-GlcNAc and 2 μg of B3GNT6 in 200 μl Labeling Buffer was prepared. For core-1 O-glycan imaging, a mixture of 20 nmol of UDP-azido-GlcNAc and 2 μg of GCNT1 in 200 μl Labeling Buffer was prepared. For HS imaging, a mixture of 20 nmol of UDP-azido-GlcNAc, 50 nmol of UDP-GlcA and 4 μg of EXT1/2 in 200 μl Labeling Buffer was prepared. For HA imaging, a mixture of 20 nmol of UDP-azido-GlcNAc, 50 nmol of UDP-GlcA and 10 μg of HAS in 200 μl Labeling Buffer was prepared. The mixtures were then applied to the wells and incubated at 37 °C for 1 to 2 hours, or at room temperature for 16 hours.

### Conjugation of the clickable carbohydrates to biotin and fluorescent dye

After the incorporation of the azido carbohydrates to cells, a biotin moiety was conjugated to the carbohydrates via copper mediated click chemistry. For each reaction, 20 nmol of Cu^2+^, 10 nmol of Biotinylated Alkyne and 200 nmol of ascorbic acid were combined in less than 10 μl of volume in a test tube and incubated at room temperature for 1 minute to let the Cu^2+^ reduce to Cu^+^. The mixture was then diluted with 200 μl of 25 mM Tris, 150 mM NaCl at pH 7.5 and then applied to a single well of cells in a 24-well plate for 30 minutes at room temperature. The reaction solution was then removed and the cells were washed thoroughly with PBS. A fluorescent dye mix of streptavidin-Alexa Fuor 555 or streptavidin-Alexa Fluor 488 at 10 μ/mL and DAPI at 10 μM in 200 μL of PBS was then applied to the cells for 15 minutes. The cells were then washed thoroughly with PBS and stored in PBS for imaging. For staining with Cy5, Biotinylated Alkyne was replaced with Cy5-Alkyne.

### Imaging of the fluorescent labeled cells

All images were captured on an AXIO Observer microscope (ZEISS) with a ZEISS Axiocam 506 mono camera and Zen 2 Pro software. Images were captured simultaneously through the channels of Alexa Fluor 555 (or Alexa Fluor 488) and DAPI. For most images, exposure time was automatically set by Set Exposure and contrast was adjusted by Best Fit. For comparison of the results of different pretreatment conditions, such as HPSE pretreatment versus Hep III pretreatment on HS staining, same exposure and contrast adjustment parameters were used. The γ ratting was not changed except where indicated.

### Cytoplasmic and nuclear extracts preparation and their pretreatment

HUVEC cells were cultured in HUVEC media (R&D system, CCM027) under 5% CO_2_. HEK293 cells were cultured in IMDM media supplemented with 5% FBS and penicillin/streptomycin. About 30x10^6 cells were scraped and collected by centrifugation at 2000x rmp for 5 minutes. The fractionation of cells was done by using NE-PER nuclear and cytoplasmic extraction reagent kit (Thermo, Cat# 78833) following the manufacturer’s instruction. In order to prevent protein degradation, protease inhibitor cocktail (Thermo, Cat# 78429) and phosphatase inhibitor cocktail (Thermo, Cat# 78426) were added to the buffers. The nuclear and cytoplasmic fractions were stored at -20 °C until it was used. For Tn antigen synthesis, 10 μl of extracts was incubated with 1 μg each of GALNT2 and GALNT3, 10 nmol of UDP-GalNAc in 25 mM Tris, 10 mM Mn^2+^,150 mM NaCl at pH 7.5 at 37°C for 1 hour. For T antigen synthesis, 10 μl of extracts was incubated with 1 μg each of GALNT2 and GALNT3, 1 μg of C1GalT1, 10 nmol of UDP-GalNAc, 10 nmol of UDP-Gal in 25 mM Tris, 10 mM Mn^2+^,150 mM NaCl at pH 7.5 at 37°C for 1 hour. For OGA treatment, 10 μl of extracts was mixed with 10 μl 0.1 M MES at pH 5.0 and 1 μg of OGA and incubated at 37°C for 30 minutes.

### SDS-PAGE and membrane blotting

SDS-PAGE and blotting were performed as described previously (Wu, Z.L., Huang, X., et al. 2015, Wu, Z.L., Huang, X., et al. 2016a).

## Acknowledgement

We would like to thank Dr. Vassili Kalabokis for his initiative support on this project; Dr. Fernando Bazan for his insightful discussion; Anuratha Elayaperumal for her help on fluorescent imaging; Dr. Timothy Manning and Dr. Diane Wotta for critical reading of the manuscriptTable I. Enzymes used in this study All are recombinant human enzymes except indicated otherwise.

## References

Abdi R, Moore R, Sakai S, Donnelly CB, Mounayar M, Sackstein R. 2015. HCELL Expression on Murine MSC Licenses Pancreatotropism and Confers Durable Reversal of Autoimmune Diabetes in NOD Mice. Stem cells, 33:1523-1531.

Ambrosi M, Cameron NR, Davis BG. 2005. Lectins: tools for the molecular understanding of the glycocode. Organic & biomolecular chemistry, 3:1593-1608.

Aruffo A, Stamenkovic I, Melnick M, Underhill CB, Seed B. 1990. CD44 is the principal cell surface receptor for hyaluronate. Cell, 61:1303-1313.

Baskin JM, Dehnert KW, Laughlin ST, Amacher SL, Bertozzi CR. 2010. Visualizing enveloping layer glycans during zebrafish early embryogenesis. Proceedings of the National Academy of Sciences of the United States of America, 107:10360-10365.

Bertozzi CR, Freeze HH, Varki A, Esko JD. 2009. Glycans in Biotechnology and the Pharmaceutical Industry. In: Varki A, Cummings RD, Esko JD, Freeze HH, Stanley P, Bertozzi CR, Hart GW, Etzler ME editors. Essentials of Glycobiology. Cold Spring Harbor (NY).

Boeggeman E, Ramakrishnan B, Kilgore C, Khidekel N, Hsieh-Wilson LC, Simpson JT, Qasba PK. 2007. Direct identification of nonreducing GlcNAc residues on N-glycans of glycoproteins using a novel chemoenzymatic method. Bioconjugate chemistry, 18:806-814.

Bond MR, Hanover JA. 2013. O-GlcNAc cycling: a link between metabolism and chronic disease. Annual review of nutrition, 33:205-229.

Brockhausen I, Schachter H, Stanley P. 2009. O-GalNAc Glycans. In: Varki A, Cummings RD, Esko JD, Freeze HH, Stanley P, Bertozzi CR, Hart GW, Etzler ME editors. Essentials of Glycobiology. Cold Spring Harbor (NY).

Chaubard JL, Krishnamurthy C, Yi W, Smith DF, Hsieh-Wilson LC. 2012. Chemoenzymatic probes for detecting and imaging fucose-alpha(1-2)-galactose glycan biomarkers. Journal of the American Chemical Society, 134:4489-4492.

Clark PM, Dweck JF, Mason DE, Hart CR, Buck SB, Peters EC, Agnew BJ, Hsieh-Wilson LC. 2008. Direct in-gel fluorescence detection and cellular imaging of O-GlcNAc-modified proteins. Journal of the American Chemical Society, 130:11576-11577.

Codelli JA, Baskin JM, Agard NJ, Bertozzi CR. 2008. Second-generation difluorinated cyclooctynes for copper-free click chemistry. Journal of the American Chemical Society, 130:11486-11493.

Crampton SP, Davis J, Hughes CC. 2007. Isolation of human umbilical vein endothelial cells (HUVEC). Journal of visualized experiments: JoVE:183.

Cummings RD, Etzler ME. 2009. Antibodies and Lectins in Glycan Analysis. In: Varki A, Cummings RD, Esko JD, Freeze HH, Stanley P, Bertozzi CR, Hart GW, Etzler ME editors. Essentials of Glycobiology. Cold Spring Harbor (NY).

Dalziel M, Crispin M, Scanlan CN, Zitzmann N, Dwek RA. 2014. Emerging principles for the therapeutic exploitation of glycosylation. Science, 343:1235681.

DeAngelis PL, Oatman LC, Gay DF. 2003. Rapid chemoenzymatic synthesis of monodisperse hyaluronan oligosaccharides with immobilized enzyme reactors. The Journal of biological chemistry, 278:35199-35203.

Dennis RJ, Taylor EJ, Macauley MS, Stubbs KA, Turkenburg JP, Hart SJ, Black GN, Vocadlo DJ, Davies GJ. 2006. Structure and mechanism of a bacterial beta-glucosaminidase having OGlcNAcase activity. Nature structural & molecular biology, 13:365-371.

Dykstra B, Lee J, Mortensen LJ, Yu H, Wu ZL, Lin CP, Rossi DJ, Sackstein R. 2016. Glycoengineering of E-Selectin Ligands by Intracellular versus Extracellular Fucosylation Differentially Affects Osteotropism of Human Mesenchymal Stem Cells. Stem cells, 34:2501-2511.

Fuster MM, Esko JD. 2005. The sweet and sour of cancer: glycans as novel therapeutic targets. Nature reviews. Cancer, 5:526-542.

Gerken TA, Jamison O, Perrine CL, Collette JC, Moinova H, Ravi L, Markowitz SD, Shen W, Patel H, Tabak LA. 2011. Emerging paradigms for the initiation of mucin-type protein Oglycosylation by the polypeptide GalNAc transferase family of glycosyltransferases. The Journal of biological chemistry, 286:14493-14507.

Gilgunn S, Conroy PJ, Saldova R, Rudd PM, O’Kennedy RJ. 2013. Aberrant PSA glycosylation-a sweet predictor of prostate cancer. Nat Rev Urol, 10:99-107.

Griffin ME, Hsieh-Wilson LC. 2016. Glycan Engineering for Cell and Developmental Biology. Cell chemical biology, 23:108-121.

Hart GW, Akimoto Y. 2009. The O-GlcNAc Modification. In: Varki A, Cummings RD, Esko JD, Freeze HH, Stanley P, Bertozzi CR, Hart GW, Etzler ME editors. Essentials of Glycobiology. Cold Spring Harbor (NY).

Hovingh P, Linker A. 1970. The enzymatic degradation of heparin and heparitin sulfate. 3. Purification of a heparitinase and a heparinase from flavobacteria. The Journal of biological chemistry, 245:6170-6175.

Hsu TL, Hanson SR, Kishikawa K, Wang SK, Sawa M, Wong CH. 2007. Alkynyl sugar analogs for the labeling and visualization of glycoconjugates in cells. Proceedings of the National Academy of Sciences of the United States of America, 104:2614-2619.

Iozzo RV. 2005. Basement membrane proteoglycans: from cellar to ceiling. Nature reviews. Molecular cell biology, 6:646-656.

Iwai T, Inaba N, Naundorf A, Zhang Y, Gotoh M, Iwasaki H, Kudo T, Togayachi A, Ishizuka Y, Nakanishi H, et al. 2002. Molecular cloning and characterization of a novel UDPGlcNAc:GalNAc-peptide beta1,3-N-acetylglucosaminyltransferase (beta 3Gn-T6), an enzyme synthesizing the core 3 structure of O-glycans. The Journal of biological chemistry, 277:12802-12809.

Ju T, Brewer K, D’Souza A, Cummings RD, Canfield WM. 2002. Cloning and expression of human core 1 beta1,3-galactosyltransferase. The Journal of biological chemistry, 277:178-186.

Ju T, Cummings RD. 2005. Protein glycosylation: chaperone mutation in Tn syndrome. Nature, 437:1252.

Khoury GA, Baliban RC, Floudas CA. 2011. Proteome-wide post-translational modification statistics: frequency analysis and curation of the swiss-prot database. Scientific reports, 1.

Kizuka Y, Funayama S, Shogomori H, Nakano M, Nakajima K, Oka R, Kitazume S, Yamaguchi Y, Sano M, Korekane H, et al. 2016. High-Sensitivity and Low-Toxicity Fucose Probe for Glycan Imaging and Biomarker Discovery. Cell chemical biology, 23:782-792.

Lairson LL, Henrissat B, Davies GJ, Withers SG. 2008. Glycosyltransferases: structures, functions, and mechanisms. Annual review of biochemistry, 77:521-555.

Mao Y, Huang Y, Buczek-Thomas JA, Ethen CM, Nugent MA, Wu ZL, Zaia J. 2014. A liquid chromatography-mass spectrometry-based approach to characterize the substrate specificity of mammalian heparanase. The Journal of biological chemistry, 289:34141-34151.

Mbua NE, Li X, Flanagan-Steet HR, Meng L, Aoki K, Moremen KW, Wolfert MA, Steet R, Boons GJ. 2013. Selective exo-enzymatic labeling of N-glycans on the surface of living cells by recombinant ST6Gal I. Angewandte Chemie, 52:13012-13015.

Moremen KW, Tiemeyer M, Nairn AV. 2012. Vertebrate protein glycosylation: diversity, synthesis and function. Nature reviews. Molecular cell biology, 13:448-462.

Natsuka S, Gersten KM, Zenita K, Kannagi R, Lowe JB. 1994. Molecular cloning of a cDNA encoding a novel human leukocyte alpha-1,3-fucosyltransferase capable of synthesizing the sialyl Lewis x determinant. The Journal of biological chemistry, 269:20806.

Pang PC, Chiu PC, Lee CL, Chang LY, Panico M, Morris HR, Haslam SM, Khoo KH, Clark GF, Yeung WS, et al. 2011. Human sperm binding is mediated by the sialyl-Lewis(x) oligosaccharide on the zona pellucida. Science, 333:1761-1764.

Pinho SS, Reis CA. 2015. Glycosylation in cancer: mechanisms and clinical implications. Nature reviews. Cancer, 15:540-555.

Posey AD, Jr., Schwab RD, Boesteanu AC, Steentoft C, Mandel U, Engels B, Stone JD, Madsen TD, Schreiber K, Haines KM, et al. 2016. Engineered CAR T Cells Targeting the Cancer-Associated Tn-Glycoform of the Membrane Mucin MUC1 Control Adenocarcinoma. Immunity, 44:1444-1454.

Pratt MR, Hang HC, Ten Hagen KG, Rarick J, Gerken TA, Tabak LA, Bertozzi CR. 2004. Deconvoluting the functions of polypeptide N-alpha-acetylgalactosaminyltransferase family members by glycopeptide substrate profiling. Chemistry & biology, 11:1009-1016.

Ramakrishnan B, Qasba PK. 2002. Structure-based design of beta 1,4-galactosyltransferase I (beta 4Gal-T1) with equally efficient N-acetylgalactosaminyltransferase activity: point mutation broadens beta 4Gal-T1 donor specificity. The Journal of biological chemistry, 277:20833-20839.

Reznikoff CA, Brankow DW, Heidelberger C. 1973. Establishment and characterization of a cloned line of C3H mouse embryo cells sensitive to postconfluence inhibition of division. Cancer research, 33:3231-3238.

Rostovtsev VV, Green LG, Fokin VV, Sharpless KB. 2002. A stepwise huisgen cycloaddition process: copper(I)-catalyzed regioselective “ligation” of azides and terminal alkynes. Angewandte Chemie, 41:2596-2599.

Sarrazin S, Lamanna WC, Esko JD. 2011. Heparan sulfate proteoglycans. Cold Spring Harbor perspectives in biology, 3.

Senay C, Lind T, Muguruma K, Tone Y, Kitagawa H, Sugahara K, Lidholt K, Lindahl U, Kusche-Gullberg M. 2000. The EXT1/EXT2 tumor suppressors: catalytic activities and role in heparan sulfate biosynthesis. EMBO reports, 1:282-286.

Slawson C, Hart GW. 2011. O-GlcNAc signalling: implications for cancer cell biology. Nature reviews. Cancer, 11:678-684.

Somers WS, Tang J, Shaw GD, Camphausen RT. 2000. Insights into the molecular basis of leukocyte tethering and rolling revealed by structures of P- and E-selectin bound to SLe(X) and PSGL-1. Cell, 103:467-479.

Sorensen T, White T, Wandall HH, Kristensen AK, Roepstorff P, Clausen H. 1995. UDP-Nacetyl-alpha-D-galactosamine:polypeptide N-acetylgalactosaminyltransferase. Identification and separation of two distinct transferase activities. The Journal of biological chemistry, 270:24166-24173.

Sterner E, Flanagan N, Gildersleeve JC. 2016. Perspectives on Anti-Glycan Antibodies Gleaned from Development of a Community Resource Database. ACS chemical biology, 11:1773-1783.

Vlodavsky I, Friedmann Y, Elkin M, Aingorn H, Atzmon R, Ishai-Michaeli R, Bitan M, Pappo O, Peretz T, Michal I, et al. 1999. Mammalian heparanase: gene cloning, expression and function in tumor progression and metastasis. Nature medicine, 5:793-802.

Wu ZL, Huang X, Burton AJ, Swift KA. 2015. Glycoprotein labeling with click chemistry (GLCC) and carbohydrate detection. Carbohydrate research, 412:1-6.

Wu ZL, Huang X, Burton AJ, Swift KA. 2016a. Probing sialoglycans on fetal bovine fetuin with azido-sugars using glycosyltransferases. Glycobiology, 26:329-334.

Wu ZL, Huang X, Ethen C, Tatge T, Pasek M, Zaia J. 2016b. Non-reducing end labeling of heparan sulfate via click chemistry and a high throughput ELISA assay for heparanase. Glycobiology.

Wu ZL, Robey MT, Tatge T, Lin C, Leymarie N, Zou Y, Zaia J. 2014. Detecting O-GlcNAc using in vitro sulfation. Glycobiology, 24:740-747.

Xu D, Esko JD. 2014. Demystifying heparan sulfate-protein interactions. Annual review of biochemistry, 83:129-157.

Yeh JC, Ong E, Fukuda M. 1999. Molecular cloning and expression of a novel beta-1, 6-Nacetylglucosaminyltransferase that forms core 2, core 4, and I branches. The Journal of biological chemistry, 274:3215-3221.

Zheng T, Jiang H, Gros M, del Amo DS, Sundaram S, Lauvau G, Marlow F, Liu Y, Stanley P, Wu P. 2011. Tracking N-acetyllactosamine on cell-surface glycans in vivo. Angewandte Chemie, 50:4113-4118.

